# Glucose-Sensing ChREBP Protein in the Pathogenesis of Diabetic Retinopathy

**DOI:** 10.1101/2024.12.04.626828

**Authors:** Christopher R. Starr, Assylbek Zhylkibayev, Oleg Gorbatyuk, Alli M. Nuotio-Antar, James Mobley, Maria B. Grant, Marina Gorbatyuk

## Abstract

Glucose-sensing ChREBP and MondoA are transcriptional factors involved in lipogenic, inflammatory, and insulin signaling pathways implicated in metabolic disorders; however, limited ocular studies have been conducted on these proteins. We aimed to investigate the potential role of ChREBP in pathogenesis of diabetic retinopathy (DR). We used diabetic human and mouse retinal cryosections analyzed by immunohistochemistry. qRT-PCR was performed to quantify gene expression. To explore the role of ChREBP in rods, we generated caChREBP^RP^ mice with constitutively active (ca) ChREBP. These mice underwent retinal function testing, followed by proteomic analysis using LC-MS. Furthermore, ARPE-19 cells were infected with lentiviral particles expressing human ChREBP (ARPE-19^ChREBP^) and subjected to global proteomics. Our results demonstrate that both proteins were expressed across the retina, although with distinct distribution patterns: MondoA was more prominently expressed in cones, while ChREBP was broadly expressed throughout the retina. Elevated expression of both proteins was observed in DR. This may have contributed to rod photoreceptor degeneration as we observed diminished scotopic ERG amplitudes detected in caChREBP^RP^ mice at P35. The retinal proteomic landscape indicated a decline in KEGG pathways associated with phototransduction, amino acid metabolism, and cell adhesion. Furthermore, rod-specific caChREBP induced TXNIP expression. Consistent with altered retinal proteomics, ARPE-19^ChREBP^ cells displayed a metabolic shift toward increased glyoxylate signaling, sugar metabolism, and lysosomal activation. Our study demonstrates that ChREBP overexpression causes significant metabolic reprograming triggering retinal functional loss in mice

## 1. Introduction

Glucose sensing Mondo family proteins (MFPs), carbohydrate response element binding protein (ChREBP), and MondoA are transcriptional modulators of lipogenic, inflammatory, and insulin signaling-associated genes that are compromised in diabetic retinopathy (DR). Both MFPs exist as heterodimers forming complexes with Max-like protein X (MLX) [1],[2]. In response to increased glucose flux into cells, both MFPs translocate to the nucleus and independently bind the MLX transcription factor, activating gene expression via the ChoRE element within promoters of target genes [3]. Example of such genes include thioredoxin-interacting protein (TXNIP) and Arrestin Domain-Containing Protein 4 (ARRDC4) [3,4]. While current research suggests their overlapping roles in the regulation of lipid metabolism and TXNIP-mediated NRLP3 activation, mounting evidence indicates their independent roles in different tissues. Thus, MondoA primarily drives lipid metabolism and insulin signaling along with control of ARRDC4-mediated glucose uptake [5,6] while ChREBP controls de novo lipogenesis and glycolysis [7,8]. ChREBP is primarily expressed in the pancreas, adipose tissue, and liver, whereas MondoA is predominantly found in skeletal muscle and immune cells [9]. For instance, in mice challenged with a high-fat diet (HFD), liver-specific ChREBP overexpression leads to elevated serum triglycerides (TG) and increased inflammation [10,11]. The muscle-specific MondoA knockout, on the other hand, shows reduced expression of genes associated with insulin signaling and fatty acid metabolism [5].

ChREBP has two known functional isoforms, α and β, each with its own promoter and transcription start site [12]. The ChREBPα protein senses glucose via the activation of two domains: a low-glucose inhibitory domain (LID) and a glucose-response activation conserved element (GRACE) domain [13]. Under low glucose conditions, the LID inhibits the GRACE domain, resulting in cytoplasmic retention of ChREBPa. However, under conditions of increased glucose flux into the cell, the intramolecular inhibition of the LID is reversed, allowing for ChREBPα translocation to the nucleus, where it transactivates target genes, including ChREBPβ. ChREBPb lacks the LID domain, making it constitutively active [12].

Previous research has thoroughly investigated the roles of both MFPs in human metabolic disorders. However, only a limited number of ocular studies have focused on ChREBP and no studies have been conducted on MondoA, indicating a gap in the knowledge of these proteins in the retina [14,15]. It has been proposed that ChREBP-mediated and normoxic HIF-1α activation may be partially responsible for neovascularization in diabetic and age-related retinopathy [15]. Furthermore, ChREBP deficiency has been shown to reduce high-glucose-induced apoptosis, migration, and tube formation in human retinal microvascular endothelial cells as well as structural and angiogenic responses in the mouse retina [14]

Therapeutic approaches targeting ChREBP and MondoA have recently been proposed to control metabolic disorders [5]. The small drug inhibitor SBI-993, which deactivates transcription factors ChREBP and MondoA, was shown to reduce liver- and muscle-produced TG, suppress TXNIP and ARRDC4, enhance insulin signaling, and improve glucose tolerance in HFD-fed mice. This underscores the key roles of these proteins in controlling lipid balance, blood glucose levels, and insulin signaling [6] These studies also suggest that both ChREBP and MondoA could be targeted therapeutically in ocular conditions. However, it remains to be determined which specific retinal diseases and cell types would benefit from this approach.

In this study, we sought to better understand the expression patterns of ChREBP and MondoA, with the aim of clarifying the potential role of ChREBP in the pathogenesis of DR. We show for the first time the retinal-specific MFP expression and increased MFP expression during DR progression. Our data suggest that ChREBP is overexpressed in the retina, which may result in photoreceptor cell deterioration through alterations in molecular events. Overall, our findings point to the critical need for a thorough examination of the role of glucose-activated proteins in DR.

## 2. Materials and Methods

### Mice

Male C57BL/6J (Strain#: 000664), Akita (Strain #:003548), db/db (BKS.Cg-Dock7m +/+ Leprdb/J; Strain#: 000642), and i-Cre (Strain#: 015850) mice were obtained from the Jackson Laboratory. The eGFP^flox/wt^-caChREBP mice have been described previously [11]. eGFP^flox/wt^-caChREBP were bred with i-Cre mice to generate rod photoreceptor (RP) caChREBP^RP^ experimental mice and eGFP^flox/wt^-caChREBP (caChREBP^flox/wt^) littermate controls. All mice were kept in a 12-hour light–dark cycle with ad libitum access to food and water. All mice were housed in the University of Alabama at Birmingham (UAB) animal facility, adhering to the guidelines set by the institutional animal care and use committee (UAB-IACUC protocol no. 22104) and the Association for Research in Vision and Ophthalmology guidelines. The animals were euthanized by carbon dioxide followed by cervical dislocation.

### Retinal explants

The eyes from C57BL/6J animals at postnatal day 8 were enucleated, and the retina was gently isolated from the eyecup and placed in Neurobasal Serumfree medium (Neurobasal-A, 10888022; Invitrogen) containing 2% B27 (0080085-SA; Invitrogen, Carlsbad, CA, USA), 1% N2 (17502-048; Invitrogen), 2mM GlutaMAX (35050038; Invitrogen), and 100 units/mL penicillin–100 lg/mL streptomycin (P4333; Sigma-Aldrich Corp., St. Louis, MO, USA). Retinal explants were maintained at 37 °C and 5% CO_2_. Mannitol (19 mM; Control) (Sigma, M4125) and D-glucose (34mM; High glucose) (Sigma, G5767) were dissolved in growth medium. Explants were cultured for 24 h and analyzed using qRT-PCR. Primers for ChREBPα (forward: 5’-CGACACTCACCCACCTCTTC-3’; reverse: 5’-TTGTTCAGCCGGATCTTGTC-3’), ChREBPβ (forward: 5’-TCTG-CAGATCGCGTGGAG-3’; reverse: 5’-CTTGTCCCGGCATAGCAAC-3’), MondoA (forward: 5’-TGCTACCTGCCACAGGAGTC-3’; reverse: 5’-GACTCAAACAGTGGCTT-GATGA-3’), and GAPDH (forward: TGACGTGCCGCCTGAAGAAA; reverse: AGTG-TAGCCCAAGATGCCCTTCAG) were used to detect Mondo Family Proteins in the retina. The same primers were applied in the study of diabetic retinas in mice with T1D and T2D diabetes.

### Infection of ARPE-19

with lentiviral particles expressing ChREBP (OriGene Co, Cat#: RC220626L2V) was conducted using 100 μl of suspension. Forty-eight hours later, the ARPE-19^ChREBP^ and control cells were harvested, and the protein extracts were prepared for the global proteomic study.

### Retinal tissue and cell homogenization and extraction for mass spectrometry

This method was broadly described in our previous study [16]. Briefly, individual retinas and ARPE-19 cells were homogenized. This method was broadly described in our previous study. After the samples were trypsinized, they were loaded onto a 1260 Infinity high-performance liquid chromatography stack (Agilent Technologies) and separated using a 75-micron i.d. × cm pulled-tip C-18 column (Jupiter C18 300 Å, 5 microns, Phenomenex). We used a Thermo Q Exactive HF-X mass spectrometer equipped with a Nanospray Flex ion source (Thermo Fisher Scientific) to conduct proteomic analysis. We then analyzed the data using the Shiny Go web resource (http://bioinformatics.sdstate.edu/go/).

### Immunochemistry

Cryopreserved eyes were sectioned 12 μm thick using a Leica CM1510S cryostat (Leica, Buffalo Grove, IL). The IHC analysis was conducted on the retinal sections using anti-ChREBP (Novus Biologicals, NB 400-135) and anti-MondoA (Invitrogen, PA5-23734) antibodies (dilution 1:200). Donkey anti-Rabbit IgG and Alexa Fluor 488 (Invitrogen, A21206) were employed as secondary antibodies. Imaging analysis was conducted using a BZ-X800 fluorescence microscope (Keyence, Itasca, IL).

### Immunoblotting

Mouse retinas were gently isolated and sonicated in a RIPA buffer supplemented with 1% Halt Protease Inhibitor and a phosphatase inhibitor cocktail (Thermo Fisher Scientific, Waltham, MA, USA, #78440). The Western blot techniques described in a previous study were employed for further analysis. Protein samples (40–60 μg) were separated by SDS-PAGE and transferred to a PVDF membrane. The anti-TXNIP (VDUP1) (mouse mAb, MBL, K0205-3 1:1000) and anti-beta actin (rabbit, Sigma Aldrich, A2066, 1:5000) antibodies were used for target protein detection. Horseradish peroxidase conjugated Goat anti-Rabbit IgG (926-80011) and goat anti-Mouse IgG (926-80010) from Li-Cor were used as a secondary antibody (1:10,000). Images of membranes were captured and analyzed using the Odyssey XF system (Li-Cor).

### Electroretinography (ERG)

At postnatal (p) day 35, mice were dark-adapted overnight, and all procedures were conducted in a dark room with light adaptation for photopic response using UTAS BigShot instrument (LKC Technologies, Gaithersburg, MD, USA). 2.5% phenylephrine (Paragon BioTeck, Inc., 42702–102-15, Portland, OR, USA) and Gonak (2.5% sterile hypromellose ophthalmic demulcent solution, Alcon, Lake Forest, CA, USA) were applied to the cornea for dilation and to moisturize the ocular surface. For scotopic ERG, mice were exposed to a series of 5 flashes at various intensities: 0.025 cds/m^2^ (−20 dB), 2.5 cds/m^2^ (0dB), 25 cds/m^2^ (10dB), and 157.74 cds/m^2^ (18dB), with 45-second intervals between flashes. Following the scotopic protocol, each mouse was light-adapted for 5 minutes under a dome background light of 25 cds/m^2^. The photopic protocol involved a series of 15 flashes with 1-second intervals between each flash, at intensities of 25 cds/m^2^, and 79 cds/m^2^. Obtained results were analyzed using the LKC EM software (LKC Technologies, Gaithersburg, MD, USA).

### RNAscope

RNAscope was performed according to the manufacturer’s (Advanced Cell Diagnostics, Newark, CA) recommendations. ChREBP was analyzed using a probe specific to its mRNA (ACDbio Cat No. 558141-C3).

### Statistical analysis

A Student two-tailed paired t-test and two-way ANOVA were used to compare the two groups using Graphpad Prism 10 software. All statistical data were expressed as mean ± SE. P < 0.05 was considered significant.

## 3. Results

### 3.1. Mondo family proteins, ChREBP and MondoA, are upregulated in human and mouse diabetic retinas

We first sought to localize ChREBP and MondoA in the retina under normal and diabetic conditions. Control and diabetic human retinas were stained with antibodies against ChREBP and MondoA and examined via confocal microscopy (Figure 1A & 1B).

**Figure 1.**
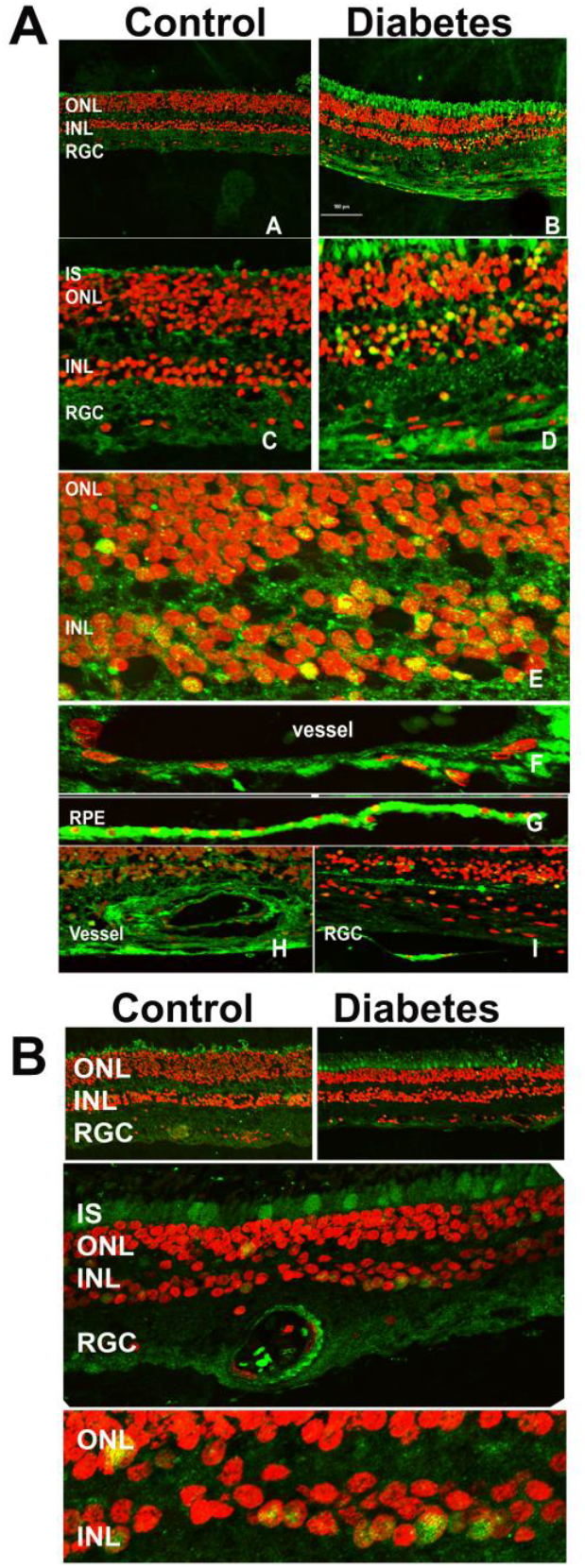
Immunohistochemical analysis of human control and diabetic retina. **A**: Retinas were subjected to treatment with anti-ChREBP antibody. The ChREBP immuno-reactivity in human control (a &c) and diabetic (b & d) retinas. In the diabetic retina, robust staining of ChREBP (green) was detected in the photoreceptors, the cells of the INL (e), endothelial cells (f & h), and retinal pigment epithelial cells (g). Co-localization of ChREBP in the nuclei (red) of retinal cells is indicated in yellow. **B**: The MondoA immunoreactivity in human control (a) and diabetic (b) retinas is shown in green. In the diabetic retina, robust staining was detected in cones (c) and the cells of the INL (d). Localization of MondoA in the nuclei is shown in yellow (d).

Immunohistochemical analysis (IHC) revealed extensive ChREBP expression across the retina (Figure 1A), with a notably stronger signal in the diabetic retinas compared to the controls. Specifically, ChREBP was observed in photoreceptors, including the outer nuclear layer (ONL) and inner segments (IS) of photoreceptors, as well as in the inner nuclear layer (INL), where it demonstrated strong immunoreactivity. A key observation was the subcellular localization of ChREBP. In diabetic retinas, ChREBP was frequently detected in nuclei, in contrast with the control retinas, where nuclear localization was significantly less pronounced. Additionally, ChREBP expression was detected in both retinal epithelial (RPE) cells (Figure 1A sub-panel G) and endothelial (HREC) cells of blood vessels (Figure 1A sub-panels F & H). These findings suggest that ChREBP is broadly expressed in the neuronal, epithelial, and endothelial cells of the retina. Moreover, its expression is enhanced in diabetic conditions, where it also shows increased nuclear translocation, indicating a potential role in mediating diabetes-induced retinal changes. Meanwhile, we observed that the expression profile of MondoA differed from that of ChREBP. MondoA expression was more prominent in cone photoreceptors than in rods. Like ChREBP, MondoA immunoreactivity was also detected in individual nuclei within the ONL and the INL.

In mouse diabetic retinas, RNAscope analysis showed elevated ChREBP mRNA levels compared to the controls (Figure 2A).

**Figure 2.**
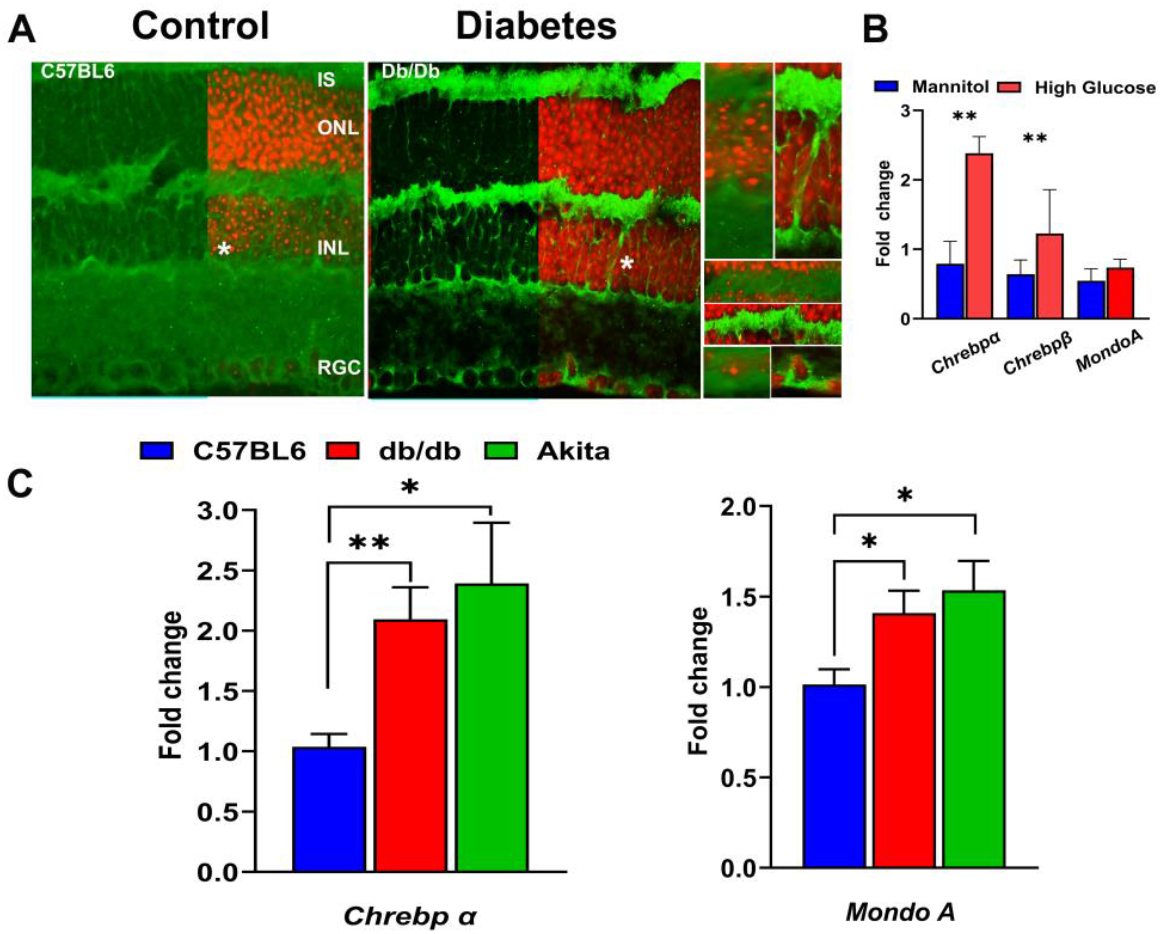
Expression of Mondo family proteins in hyperglycemic retinas. **A:** The RNAscope technique revealed enhanced ChREBP mRNA expression (green) in 12-week-old db/db retinas. The retinal ganglion cells (RGCs), Müller cells (highlighted in right inserts), and photoreceptors are responsive to hyperglycemia, showing increased ChREBP expression. **B:** Retinal explants were cultured in a medium supplemented with either high glucose or an equimolar concentration of mannitol (control) for 24 hours. High glucose, but not mannitol, culture conditions resulted in significant increases in both Chrebpα and Chrebpβ mRNAs, indicating that high glucose stimulates ChREBP expression ex vivo. **C:** To confirm ex vivo findings, diabetic retinas were isolated for qRT-PCR analysis to evaluate ChREBP and MondoA gene expression. The qRT-PCR results show that retinas from 12-week-old db/db and Akita mice exhibit increased expression of both Chrebp and Mon-doA mRNAs. Statistical significance is indicated as p*<0.05 and p**<0.01 (n=4 per group).

In 12-weeks-old db/db mice, ChREBP mRNA was detected in photoreceptors, Müller cells, and cells in the INL and retinal ganglion cells (RGCs). Furthermore, to assess whether diabetic conditions stimulated ChREBP expression in the retina, retinal explants were exposed to high-glucose media. Real-time PCR analysis revealed a marked upregulation of both ChREBPa and b isoforms due to high glucose culture conditions (Figure 2B). However, MondoA expression showed no significant response to this treatment (Figure 2B). In contrast, in vivo, both the db/db and Akita mouse retinas manifested elevated expression of ChREBP and MondoA mRNA compared to wild-type retinas (Figure 2C) although the increase in ChREBP expression in both diabetic retinas was more pronounced compared to MondoA expression. Altogether, the results presented thus far demonstrate that both Mondo family proteins are expressed in the neuronal retinas of humans and mice, with some differences in the distribution of immunoreactivity between rods and cones. Hyperglycemic conditions further stimulate the expression of ChREBP and MondoA in the retinas.

### 3.2. Overexpression of ChREBP in photoreceptors results in diminished scotopic ERG responses

To further investigate the role of ChREBP in rod photoreceptors and mimic the increased ChREBP expression observed in diabetic retina, we generated mice with constitutively active (ca) ChREBP expression in rods (Figure 3A).

**Figure 3.**
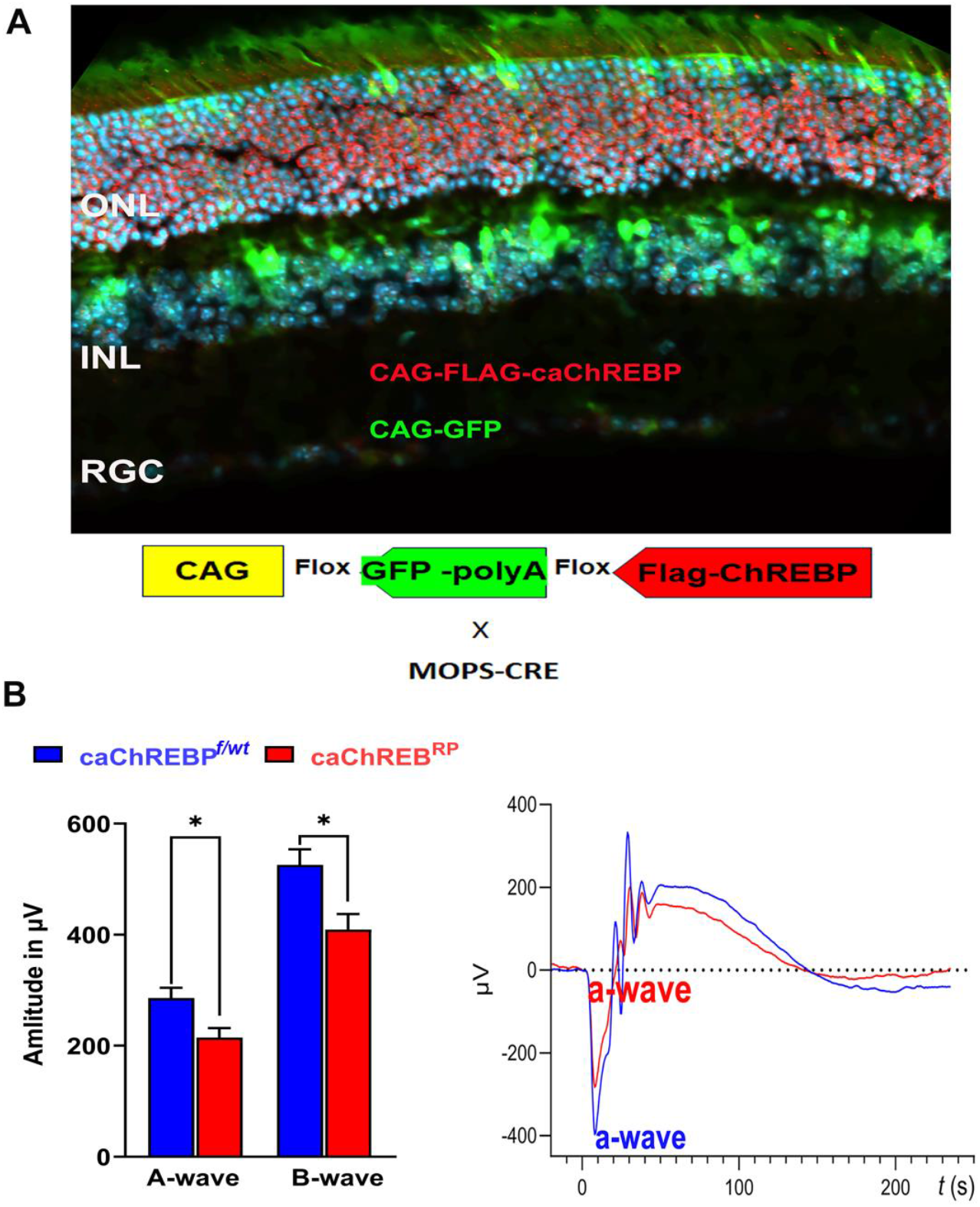
Transgenic expression of constitutively active ChREBP in rods leads to vision loss. **A:** i-Cre-mediated recombination in the rod photoreceptors of caChREBPRP transgenic mice resulted in deletion of eGFP and expression of FLAG-tagged caChREBP, detected by anti-FLAG antibody, shown in red. The nuclei are shown in blue. **B:** Expression of caChREBP in rod photoreceptors leads to reduction in the scotopic a- and b-wave amplitudes in caChREBP^RP^ versus control (eGFP^flox/wt^-caChREBP or CghREBP^f/wt^) mice at postnatal day 35. Representative waveforms are shown on the right. p*<0.05, n=6 per group.

eGFPflox/wt-caChREBP mice have been described previously and, in the absence of Cre recombinase, ubiquitously express the enhanced green fluorescent protein (eGFP), the genetic sequence for which is located between two loxP sites and which is immediately upstream of a sequence for N-terminal FLAG-tagged caChREBP [11]. Excision of the eGFP sequence and subsequent expression of FLAG-tagged caChREBP occurs following Cre recombination. I-Cre mice express Cre recombinase under the control of the short mouse rod opsin (MOPS) promoter, ensuring caChREBP expression, specifically in rod photoreceptors (RP). As expected, the caChREBP^RP^ retinas exhibited GFP expression in cells lacking Cre recombination (such as cone photoreceptors or cells of the INL), while the FLAG-tagged caChREBP protein, detected with anti-FLAG antibody and shown in red, was observed in rod photoreceptors.

In the p35 caChREBP^RP^ mice, ERG analysis revealed a reduction in scotopic a- and b-wave amplitudes compared to eGFP^flox/wt^-caChREBP (ChREBP^f/wt^) (Figure 3B). No significant loss of photopic ERG amplitude was observed in caChREBP^RP^ mice at this point. These data suggest that the ChREBP upregulation observed in diabetic retinas could contribute to the loss of retinal function in diabetic mice. Therefore, we became interested in ChREBP-mediated cellular signaling and performed a study of global proteomics with caChREBP^RP^ retinas.

### 3.3. Overexpression of ChREBP in photoreceptors alters retinal proteomics

The constitutive expression of ChREBP in rod photoreceptors of caChREBP^RP^ mice lead to substantial changes in the protein landscape within the retina, affecting several key metabolic, structural, and signaling pathways. Over 40 proteins were differentially expressed (p < 0.05) in the caChREBP-retinas, both upregulated and downregulated. The heat map of the top-modified proteins is shown in Figure 4A.

**Figure 4.**
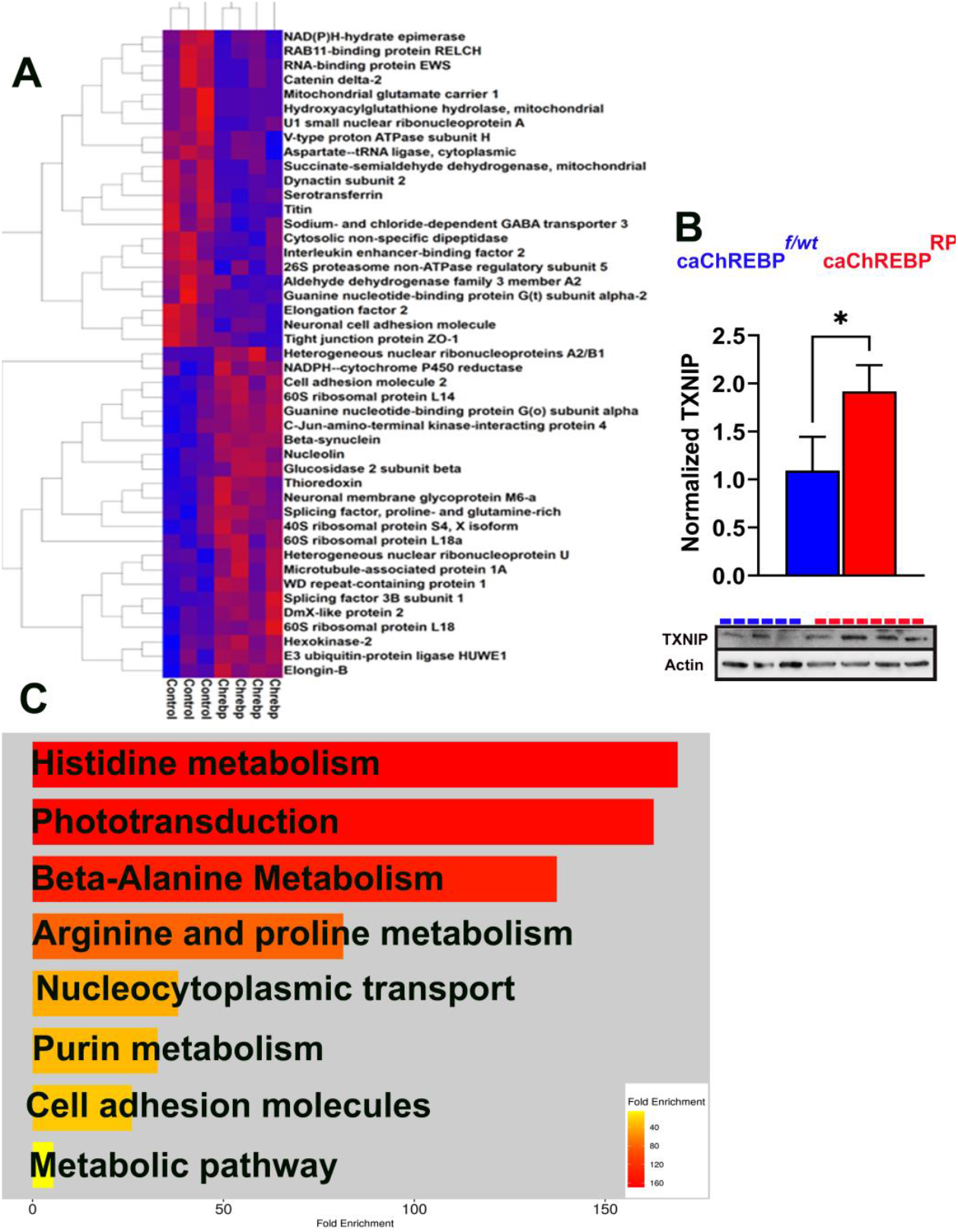
Increased ChREBP activity in rods alters retinal proteomics and KEGG signaling in caChREBP^RP^ mice. **A:** Heatmap showing the major altered proteins in caChREBP^RP^ compared with eGFP^flox/wt^-caChREBP control (Control) retinas (n=3-4 per group), highlighting significant changes in retinal protein expression.**B:** Expression of constitutively active ChREBP (caChREBP) in rods leads to an increase in TXNIP protein levels at postnatal day 35 (P35). Statistical significance is indicated by p*<0.05, with n=3-4 per group. **C:** The major downregulated KEGG signaling pathways in caChREBP-expressing retinas are identified, showing significant pathway alterations due to sustained ChREBP activity in rods.

The list of upregulated proteins included hexokinase 2, glucosidase 2, splicing factor 3B, and NADPH-cytochrome P450 reductase. The list of downregulated proteins contained tight junction protein ZO1, marker of tight junction, PDE6α, GNAT2, and Succinate-semialdehyde dehydrogenase mitochondrial (Figure S1). Figure S2 presented an analysis of ingenuity pathways modified by upstream regulators ChREBP (MLXIPL) which had the highest Z-score activation as an upstream regulator in this experiment

Given that following G6P allosteric binding, ChREBP translocates to the nucleus and forms a complex with MLX to bind the carbohydrate response element (ChoRE) in the target gene promoters, we examined its downstream signaling as this activation is central to ChREBP’s role in metabolic gene regulation, especially in conditions of high glucose. TXNIP is a well-known ChREBP target gene containing the ChoRE element [3]. This protein is also significantly implicated in DR, as it inhibits a major antioxidant protein thioredoxin. Elevated TXNIP in diabetic retina is thought to contribute to cellular stress and disease progression. In our study, TXNIP expression increased by about 50% in caChREB-PRP retinas, consistent with ChREBP’s role in promoting TXNIP-induced cell death (Figure 4B) [17].

Using ShinyGO 8.0, we then analyzed a list of downregulated KEGG signaling pathways in caChREBP^RP^ retinas (Figure 4C). This analysis revealed significant downregulation in pathways related to histidine, alanine, arginine, and purine metabolism. Consistent with our previous findings on retinal physiology, the phototransduction pathway was identified as one of the primary downregulated KEGG pathways, indicating a potential decline in retinal function. These findings indicate that the upregulation of proteins involved in glucose metabolism, glycoprotein biosynthesis, and oxidative stress, together with protein changes affecting the visual cycle, mitochondrial energy, and amino acid metabolism, underscores ChREBP’s influence on retinal metabolism and structure.

### 3.4. Overexpression of human ChREBP in ARPE-19 cells causes metabolic reprogramming and alteres cellular signaling

Hyperglycemia also affects RPE cell health as well [18-20]. Therefore, knowing that ChREBP is expressed in RPE [15], we next infected the ARPE-19 cells with lentiviral particles expressing ChREBP for 48 h. After harvesting, cells were subjected to preparation of protein extracts and global proteome analysis by LC-MS. Overall, 36 proteins were differentially expressed in the treated RPE group and ChREBP was one of them (Figure S3). Figure 5 depicts the heat map of the top differentially expressed proteins with p < 0.05. This map indicates that hexokinase-1, LAMP1, L AMP2, and mitochondrial Acetyl-coA acyltransferase were upregulated, while the large ribosomal uL23 and 26proteosome non-ATPase regulatory subunits were downregulated.

**Figure 5.**
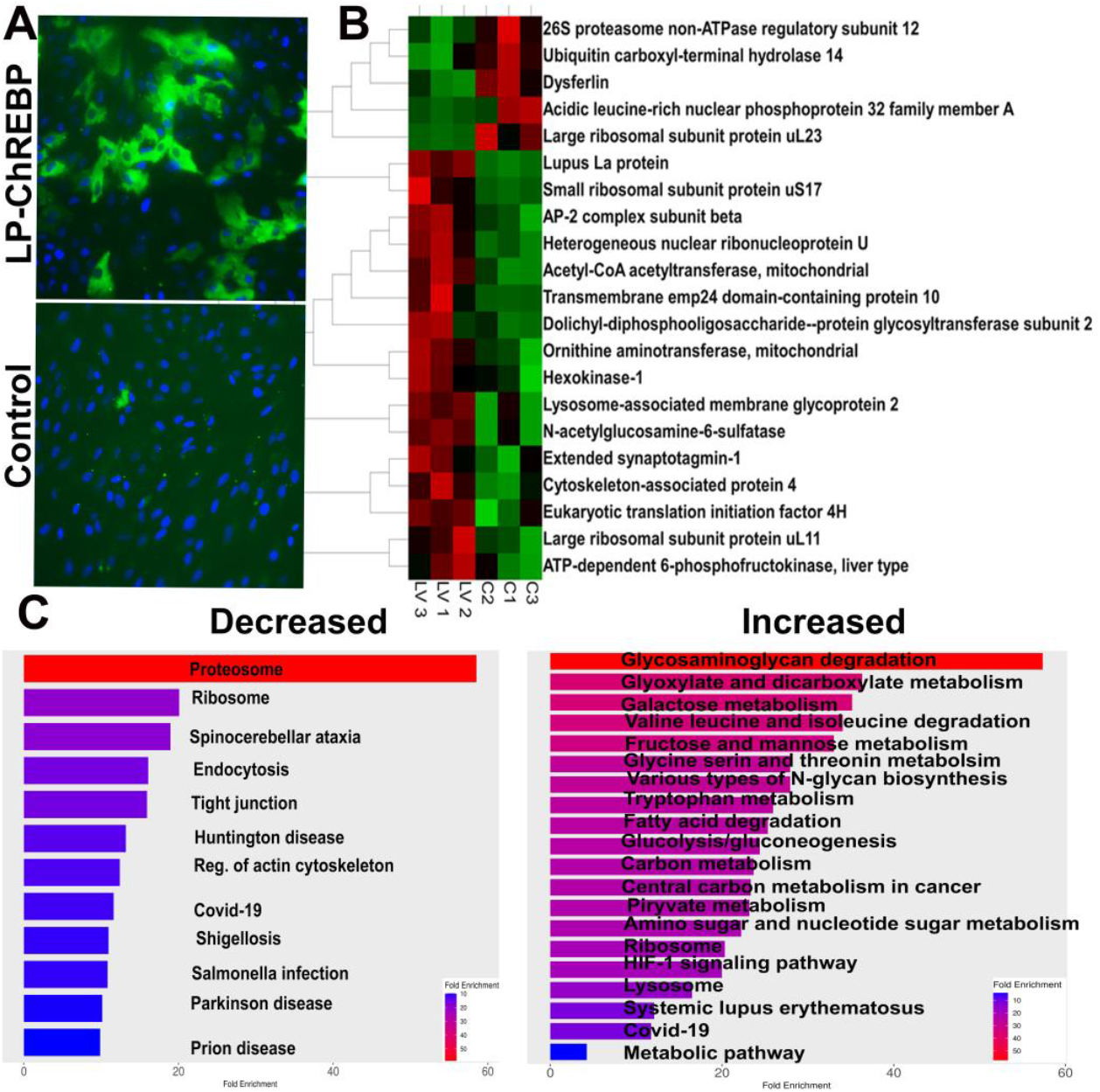
Overexpression of ChREBP in ARPE-19 Cells. **A:** Images of ARPE-19 cells overexpressing human ChREBP and control cells treated with empty virus. Direct fluorescence emitted by GFP indicates successful infection and ChREBP expression. **B:** The heatmap displays the major altered proteins in ARPE-19 cells with sustained ChREBP expression (N=3-4 per group), highlighting significant protein expression changes due to ChREBP overexpression. **C:** Results from the proteomic analysis were analyzed using the Shiny GO program to generate diagrams of altered KEGG pathways. Both decreased and increased pathways are shown, reflecting the impact of sustained ChREBP expression on cellular signaling networks in ARPE-19 cells.

Analysis of KEGG signaling pathways indicates that glyoxylate and dicarboxylate metabolism, sugar (including galactose, fructose, and mannose) metabolism, and glycolysis itself increased in ARPE-19 cells overexpressing ChREBP. These changes aligned with the activation of the lysosome pathway. Meanwhile, as observed for the retina with ChREBP overexpression in photoreceptors, tight junction signaling was diminished following treatment, along with a reduction in proteasome signaling. Further analysis of altered molecular function in treated cells demonstrated an increase in acetyl-CoA acetyl-transferase activity, while TORC2 complex binding activity was downregulated (Figure 6).

**Figure 6.**
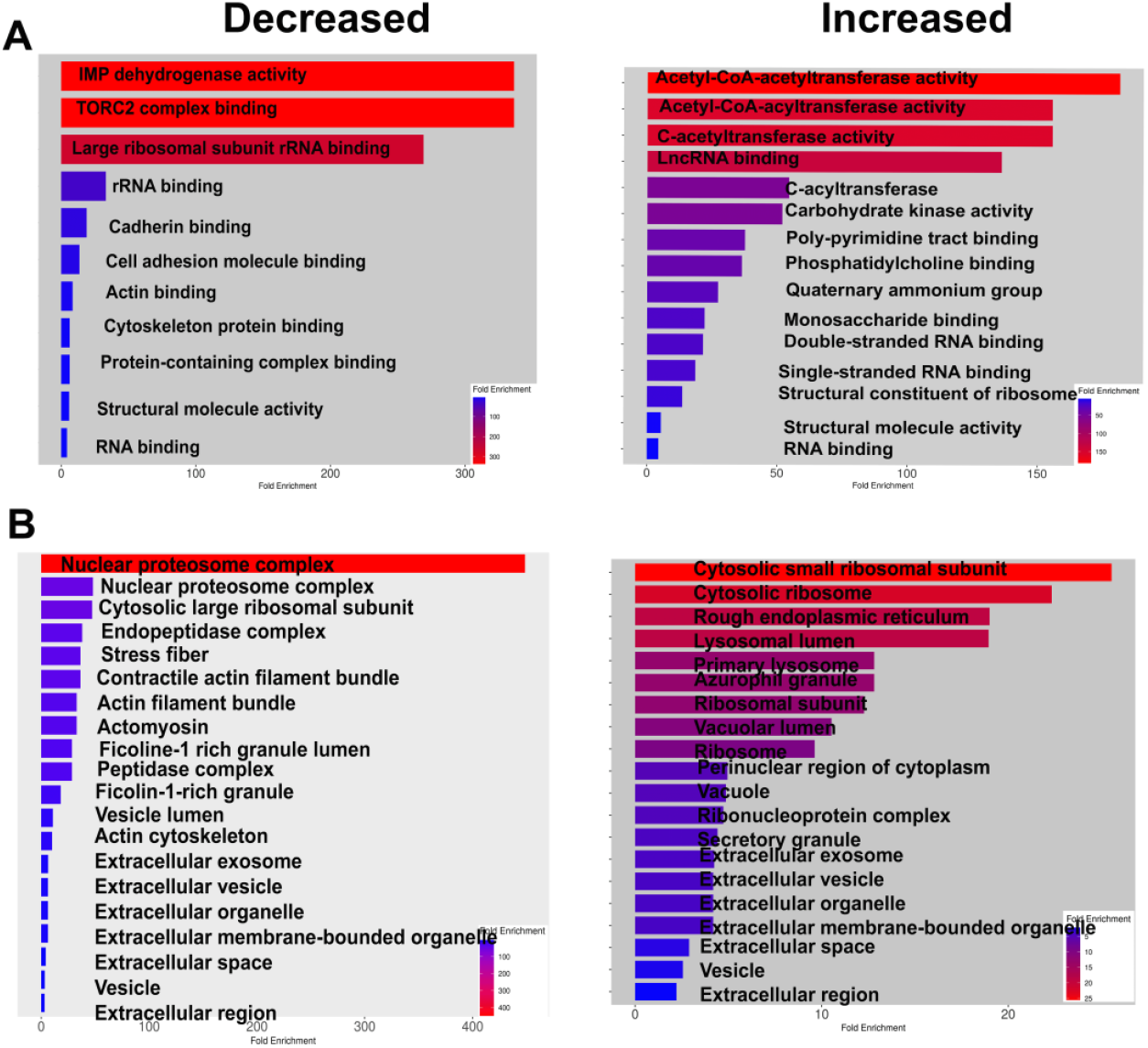
ChREBP Overexpression Alters Molecular Function and Affects Cellular Components of ARPE-19 Cells. **A:** The molecular functions that are decreased or increased due to ChREBP overexpression in ARPE-19 cells are shown, highlighting the functional alterations induced by sustained ChREBP expression. **B:** The cellular components affected by ChREBP expression in ARPE-19 cells are presented, demonstrating how ChREBP over-expression leads to changes in the cellular architecture and composition (n=3-4 per group).

Consistent with observations in the with ChREBP^RP^ retina, the cell adhesion complex was reduced upon ChREBP overexpression, including cadherin binding. The function of major cellular components, such as the nuclear proteasome and proteosome complexes, declined. In addition, ChREBP overexpression enhanced the formation of a cytosolic small ribosomal complex and lysosomes (Figure 6).

## 4. Discussion

The role of the MondoA family of proteins has been studied in various metabolic diseases, including cancer and diabetes. With few exceptions in the literature highlighting the role of ChREBP in endothelial cells and RPE, no studies have been conducted on healthy or diseased retinal tissues, indicating a gap in our knowledge of DR pathogenesis [14,15]. To the best of our knowledge, this is the first study to demonstrate the expression of ChREBP and MondoA in human retinas. Furthermore, we also investigated ChREBP and MondoA expression during the progression of DR and found that both mRNAs were elevated in T1D and T2D mouse retinas. Using transgenic mice, we replicated the increased ChREBP expression in retinas observed under diabetic conditions in the retinas and found that ChREBP overexpression in rod photoreceptors appears to be harmful, leading to retinal functional loss associated with alteration of metabolic pathways. Finally, not only photoreceptor cells but also RPE cells overexpressing ChREBP manifest metabolic reprogramming and compromised molecular function, highlighting the potential role of ChREBP in diabetic RPE.

The retina is a highly metabolic tissue, making it particularly vulnerable to metabolic disturbances, such as DR. Indeed, the retinal glucose level rises upon sustained hyperglycemia controlling expression of glucose-regulated proteins and leading to unscheduled glycolysis and activation of polyol pathway, hexosamine, PKC and AGE pathways [20-22]. Both glucose-sensing ChREBP and MondoA proteins are expressed in the retina, although their expression patterns differ. ChREBP is expressed in photoreceptors, endothelial cells (vessels), epithelial cells (RPE), and other neuronal cells (INL). In contrast, MondoA expression appears to be more robust in cone photoreceptors. These results are in agreement with the depository database “The human protein atlas,” suggesting its expression in cones is the highest in the entire human body (https://www.proteinat-las.org/ENSG00000175727-MLXIP/single=cell). Diabetes induces an increase in ChREBP and MondoA mRNA levels in the retina. High glucose regulates the expression of both proteins. In cases of hyperglycemia, following G6P-promoted allosteric conformational changes, both MFPs translocate to the nucleus and independently bind the MLX transcription factor, activating the expression of genes via the ChoRE element within the promoters of target genes, such as TXNIP. In line with this, we observed strong ChREBP immunoreactivity in the nuclei of photoreceptors and other neurons whose nuclei could be found in the retina’s inner nuclear layer upon hyperglycemia. Furthermore, overexpression of ChREBP in photoreceptors leads to an increase in TXNIP protein, suggesting a mechanism through which DR directly affects cell viability.

Transgenic caChREBP overexpression results in a decline in retinal function. Both a- and b-wave scotopic ERG amplitudes are diminished. Ongoing research on retinal dysfunction has reported reduced cone sensitivity [23], delayed activation of cone phototransduction cascade [24], selective loss of S cones [25,26], glial abnormalities, and thinning of both the nerve fiber and the RGC layer in individuals with DR [27-29] and animal models of DR [21,30-32]. Our data point to a mechanism through which diabetes, associated with sustained ChREBP activity and increased expression of ChoRE element-containing target genes such as TXNIP in the retina, may result in loss of vision and an altered proteomic landscape of the diabetic retina

Indeed, ChREBP overexpression results in an altered protein profile in p35 caChREBP^RP^ retinas. These changes were mostly associated with elevated glucose metabolism. For example, hexokinase-2 initiates glycolysis. Glucosidase 2, a key enzyme for glycoprotein biosynthesis, processes glycoproteins. SF3 is a part of the splicing machinery complex and regulates gene expression. Meanwhile, the caChREBP^RP^ retina manifests compromised cell-to-cell adhesion and blood-retinal barrier integrity, as ZO1 protein levels decline. In line with this, a reduction in proteins responsible for mitochondrial energy metabolism, such as the glutamate carrier and sodium- and chloride-dependent GABA transporter 3, was observed. The compromised expression of these proteins, along with reduced rod-specific PDE6α, is most likely associated with a decline in phototransduction, as mice deficient in PDE6α manifest dramatic vision loss [33].

Another change observed in the diabetic retina included compromised histidine and alanine metabolism. In addition to their roles as amino acids essential for protein synthesis, histidine can be converted to histamine, a neurotransmitter and immune modulator whose deficiency may contribute to neurological problems. A lack of alanine in the retina may further disrupt cellular function too, as alanine plays a critical role in energy metabolism, serving as a key substrate in the glucose-alanine cycle. Disruption of this cycle could lead to impaired energy balance and accumulation of toxic byproducts, potentially contributing to retinal dysfunction. In line with these findings, ChREBP overexpression in RPE cells alters cellular metabolism as well. Indeed, RPE cells have several unique metabolic features. For example, they manifest the high amount of lipid processing necessary for the visual cycle as they regenerate visual pigments. RPE phagocytose photoreceptor outer segments and routinely digest photoreceptor outer segments, a highly demanding process that involves extensive lipid and protein degradation. RPE cells contain robust antioxidant systems, including elevated levels of glutathione, superoxide dismutase, and catalase, because of their adaptation to high metabolic activity, constant light exposure, and oxygen consumption. Finally, the RPE cells rely on both oxidative phosphorylation and glycolysis to meet its energy demands, and their metabolism strongly depends on lactate. Therefore, the observed metabolic reprogramming disturbs homeostasis.

Glyoxylate has been proposed as a new metabolite marker of T2D [34]. A significant increase in glyoxylate levels in plasma was observed three years prior to the diagnosis of diabetes in humans and in db/db mice. Moreover, it has been proposed that elevated glyoxylate may contribute to an increase in advanced glycation end products [34]. RPE cells overexpressing ChREBP manifested an increase in glyoxylate and dicarboxylate metabolism, supporting that sustained elevation of ChREBP during diabetes may enhance the conversion of enhanced glyoxylate to advanced glycation end produces (AGEs). In support of this hypothesis, the proteasome system is compromised in treated ARPE-19 cells, which could create conditions favorable for AGE production [35].

The decline in the function of the nuclear proteasome and proteosome complexes could indicate impaired protein degradation, which may lead to the accumulation of damaged or unnecessary proteins. Meanwhile, ChREBP overexpression enhances lysosomal activity, to manage the excess of generated proteins or to compensate for the reduced protein degradation capacity under conditions influenced by ChREBP overexpression. In agreement with0df this, the LAMP1 and LAMP2 proteins, as well as the HIF-1 signaling for which lysosomes act as a degradation organelle [36], were upregulated. Moreover, the highest increase in glycosaminoglycan degradation supports this hypothesis as well; GAGs, a long-chain of carbohydrates, are degraded by lysosomes and a major component of extracellular matrix and the cell surface, activate cellular components of the treated cells [37].

## 5. Conclusions

In conclusion, this study is the first to demonstrate that Mondo family proteins, ChREBP and MondoA, are expressed throughout the retina and upregulated in humans with diabetic retinopathy and in murine models. This overexpression reprograms cellular metabolism, leading to vision loss. Therefore, these findings highlight a pivotal role for elevated ChREBP in DR and emphasize the therapeutic potential of targeting ChREBP and MondoA, paving the way for further research into novel treatments.

## Supporting information

https://zenodo.org/records/14277302

## Supplementary Materials

The following supporting information can be downloaded at: www.mdpi.com/xxx/s1, Figure S1: Quantitative spectrum counts for the proteomic analysis of caChREBP^RP^ retinas; Figure S2: Ingenuity pathways significantly changed by ChREBP overexpression in rods of caChREBP^RP^ mice. Upstream regulator MLXIPL/ChREBP had the highest positive Z score activating transcriptional program; Figure S3: Quantitative spectrum counts for the proteomic analysis of the ARPE-19 cells overexpressing ChREBP.

## Author Contributions

Conceptualization, M.G.; methodology, M.G., C.S, A.Z.; software, J.M., C. S.; validation, C.S., O.G; formal analysis, C. S., M.G.; resources M.G., A. N-A.; data curation, M.G., C.S, A.Z., O.G; writing—original draft preparation, M.G., C.S; writing—review and editing, M.G, C.S., A.N-A.; visualization, M.G., O.G., C.S., A.Z., J.M; supervision, M.G.; project administration, M.G. All authors have read and agreed to the published version of the manuscript.

## Institutional Review Board Statement

The animal study protocol was approved by the University of Alabama at Birmingham - Institutional animal care and use committee (protocol no. 22104).

## Data Availability Statement

The research data is available upon request . The raw data are also available as a supplemental material.

## Acknowledgments

Normal human and diabetic retinas were a gift of Dr. Mohamed Al-Shabrawey at Oakland University.

