## Supplementary material for "Glucose-Sensing ChREBP Protein in the Pathogenesis of Diabetic Retinopathy": https://zenodo.org/records/14277302: fig. S2.docx

| **Upstream Regulator** | **Molecule Type** | **Activation z-score** | **p-value of overlap** | **Target Molecules in Dataset** |
| --- | --- | --- | --- | --- |
| GABA | chemical - endogenous mammalian | -2.5 | 0.00000285 | ACADM,ALDH5A1,ATP6V1E1,DDX39B,  GNGT1,HADH,PRKCSH,RAP1B,RPL18,RPL30,RPS18,RPS25,RPS26 |
| AR | ligand-dependent nuclear receptor | -2.4 | 0.356 | ACTR3,ALDH1A1,CEND1,PRKAR2A,TF,TJP1 |
| 8-bromo-cAMP | chemical reagent | -2.2 | 0.229 | ALDH3A2,ASS1,CRYAA/LOC102724652,EZR,MYH9 |
| FMR1 | translation regulator | -2.2 | 3.19E-17 | ATP5IF1,ATP6V1E1,CEND1,EEF2,RPL10A,RPL11,  RPL14,RPL18,RPL18A,RPL3,RPL30,RPL7A,RPL8,RPS18,RPS20,RPS25,RPS26,  RPS4Y1,SYN1,TF |
| JUN | transcription regulator | -2.2 | 0.000115 | ANXA1,CRYAA/LOC102724652,DARS1,EEF2,EZR,HK2,HNRNPA2B1,HNRNPU,  MYH9,NRCAM,TF,TXN |
| CST5 | other | -2.1 | 0.000329 | EEF1D,EZR,HNRNPA2B1,HNRNPU,NCL,RTN3,SLC1A3,TXN |
| butyric acid | chemical - endogenous mammalian | -2.1 | 0.178 | ALDH1A1,ANXA1,MYH14,NCL,PRKAR2A |
| VDR | transcription regulator | -2.0 | 0.0957 | CLIP1,HK2,TJP1,TXN |
| medroxyprogesterone acetate | chemical drug | -2.0 | 0.357 | ALDH3A2,ASS1,EZR,MYH9 |
| PML | transcription regulator | -2.0 | 0.0199 | ACADM,CACYBP,SUMO2,TXN |
| torin1 | chemical reagent | -2.0 | 4.14E-15 | ATP5PB,ATP6V1E1,EEF1D,EEF2,HK2,RPL10A,RPL11,RPL14,RPL18,RPL3,RPL30,  RPL7A,RPL8,RPS18,RPS20,RPS25,TPT1 |
| 5-fluorouracil | chemical drug | -1.8 | 4.51E-08 | ABAT,ATP5PB,EEF2,GTF2I,HNRNPA2B1,HNRNPAB,ILF2,NACA,NCL,RPL10A,RPL11,  RPL18,RPL30,RPL7A,RPS4Y1 |
| LARP1 | translation regulator | -1.6 | 1.61E-16 | EEF1D,EEF2,RPL10A,RPL11,RPL14,RPL18,RPL3,RPL30,RPL7A,RPL8,RPS18,RPS20,  RPS25,TPT1 |
| sirolimus | chemical drug | -1.6 | 0.000000045 | ACADM,ASS1,EEF2,FSCN1,HK2,HNRNPU,NCL,RPL10A,RPL18,RPL18A,RPL3,RPL30,  RPL7A,RPL8,RPS18,RPS20,RPS26,RPS4Y1,TJP1 |
| PAX3-FOXO1 | fusion gene/product | -1.4 | 0.00242 | ABAT,ANXA1,DNAJA2,EEF2,MYH9,NRCAM,RAB5C,TCP1 |
| PTEN | phosphatase | -1.4 | 0.00417 | ANXA1,CA2,CLTA,GPHN,GPM6A,GTF2I,HADH,PLEC,PLXNB2,PPM1A,RAB5A,RAB5C |
| RICTOR | other | -1.4 | 4.19E-11 | ATP5PB,ATP6V1E1,ATP6V1H,HK2,PSMD1,RPL10A,RPL11,RPL14,RPL18,RPL30,  RPL7A,RPL8,RPS18,RPS26,Rps3a1,RPS4Y1 |
| aflatoxin B1 | chemical - endogenous non-mammalian | -1.3 | 0.036 | ALDH1A1,CA2,CRYAB,EZR,NRCAM |
| TP73 | transcription regulator | -1.3 | 0.0387 | CRYAB,Elob,HAGH,LASP1,MYH9,PLEC,UBE2V1 |
| MYOD1 | transcription regulator | -1.3 | 0.00104 | ASS1,CRYAB,ENO3,HADH,RPL7A,TJP1,TTN |
| SRF | transcription regulator | -1.2 | 0.0102 | ACTR3,EZR,GPM6A,MYH9,TJP2,TTN,WDR1 |
| benzo(a)pyrene | chemical toxicant | -1.1 | 0.0133 | ANXA1,GNAO1,HK2,NBEA,TF |
| ST1926 | chemical drug | -1.0 | 0.000000757 | AP2S1,EEF2,ILF2,NACA,RPL18A,RPL8,RPS26,TPT1,TXN |
| nicotinamide-beta-riboside | chemical - endogenous mammalian | -1.0 | 0.000394 | ALDH3A2,ALDH5A1,HADH,POR |
| NAMPT | cytokine | -1.0 | 0.00464 | ALDH3A2,ALDH5A1,HADH,POR |
| tanespimycin | chemical drug | -1.0 | 0.00928 | HNRNPA2B1,HNRNPAB,RPL30,SYN1 |
| CTNNB1 | transcription regulator | -0.8 | 4.69E-10 | ACTR3,ALDH1A1,ALDH3A2,ANXA1,CTNND2,ENO3,Ewsr1,GNAO1,HK2,NRCAM,  PRKCSH,RPL10A,RPL11,RPL14,RPL18,RPL18A,RPL3,RPL30,RPL7A,RPL8,RPS18,RPS20,  RPS25,RPS26,Rps3a1,RPS4Y1,TJP1 |
| EGR2 | transcription regulator | -0.8 | 0.00924 | ASS1,NCL,Ptms,SLC1A3,TJP2,TOP2B |
| CD 437 | chemical drug | -0.7 | 0.0000414 | AP2S1,EEF2,FSCN1,ILF2,NACA,RPL11,RPL8,TXN |
| topotecan | chemical drug | -0.7 | 0.00182 | ASAP2,FUBP1,NBEA,NIPSNAP2,PPM1A,RAB10,RAB5A,SFPQ |
| AHR | ligand-dependent nuclear receptor | -0.7 | 0.0341 | ALDH3A2,ALDH5A1,CRYAB,DMXL2,Gm21596/Hmgb1,HK2,MYH9,TXN |
| ESRRG | ligand-dependent nuclear receptor | -0.7 | 0.00709 | ACADM,HADH,HK2,TTN |
| NFE2L2 | transcription regulator | -0.6 | 0.0266 | DDX39B,HK2,PSMD1,PSMD5,RPL18,Snrpa (includes others),TJP1,TXN |
| calcitriol | chemical drug | -0.5 | 0.026 | CA2,HAGH,HK2,Hmgb3,POR,TF,TJP1,TJP2,TOP2B,TXN |
| L-triiodothyronine | chemical - endogenous mammalian | -0.5 | 0.0296 | ALDH1A1,ASS1,KRT17,OPN1LW,POR,RGS9,TF,TTN |
| pidnarulex | chemical drug | -0.4 | 0.000214 | ACTR3,ALDH1A1,ASS1,RPS4Y1,TPT1 |
| ASPSCR1-TFE3 | fusion gene/product | -0.4 | 0.00037 | ATP6V1E1,CRYAB,EZR,PACSIN2,SNCB |
| TEAD1 | transcription regulator | -0.4 | 0.00408 | ACADM,ATP5PB,ETFA,HADH,TTN |
| HGF | growth factor | -0.4 | 0.0149 | CRYAB,DDX3X,HK2,KRT17,NRCAM,RAB5A,SLK,TJP1 |
| AKT1 | kinase | -0.4 | 0.0425 | ASS1,ATP6V1E1,CRYZ,EZR,Gm5938 (includes others) |
| VEGF (family) | group | -0.4 | 0.0179 | CA2,CRYAB,DDX3X,HK2,NRCAM,RGS9,SLK,TJP1 |
| HMGA1 | transcription regulator | -0.4 | 0.00663 | CNDP2,CRYAB,MYH9,RPL7A,TRIM28 |
| ciprofloxacin | chemical drug | -0.4 | 0.00000773 | ATP6V1E1,ATP6V1H,NRCAM,PLXNB2,RPL10A,RPL7A,RPS18 |
| TYA-018 | chemical reagent | -0.4 | 0.000286 | ACADM,ATP5PB,ETFA,GPHN,HADH,MYH14,TTN |
| Z-LLL-CHO | chemical - protease inhibitor | -0.4 | 0.00019 | CRYAA/LOC102724652,CRYAB,CTNND2,HNRNPA2B1,PPM1A,PSMD1,RAB10,  TJP1,TRIM28,TXN |
| GLI1 | transcription regulator | -0.3 | 0.0226 | ANXA1,EZR,HNRNPU,HUWE1,KRT17,PPM1A,RPL30,SLC1A3 |
| PD98059 | chemical - kinase inhibitor | -0.3 | 0.0121 | ACTR3,ANXA1,CA2,GTF2I,HNRNPA2B1,HNRNPAB,PLEC,TF,TXN |
| NR1I2 | ligand-dependent nuclear receptor | -0.3 | 0.0107 | ALDH1A1,ALDH3A2,POR,TF,TJP1 |
| methylprednisolone | chemical drug | -0.3 | 0.00665 | ABAT,ALDH1A1,ALDH3A2,ANXA1,ASS1,ATP5IF1,HAGH,HSD17B10,NCL,POR |
| dexamethasone | chemical drug | -0.3 | 0.000461 | ACADM,ACTR3,ALDH1A1,ANXA1,ASAP2,CA2,CHD4,CRYAA/LOC102724652,  CRYAB,ETFA,EZR,FSCN1,GTF2I,HK2,HNRNPAB,ILF2,KPNB1,KRT17,PLEC,POR,  PRKAR2A,Rps3a1,SLC1A3,SLC25A22,TF,TJP1,TPT1,TXN,WDR1 |
| APP | other | -0.2 | 0.000201 | ARF5,ATP6V1E1,CHD4,CLTA,GDI2,GNAO1,KPNB1,MADD,PDXP,PRKAR2A,  RAB5A,SNCB,SYN1,TJP1,TOP2B,TPT1,TXN |
| EPO | cytokine | -0.2 | 0.0000291 | CA2,ILF2,KPNB1,NACA,Rps3a1,RPS4Y1,TF,TJP1,TPT1,WDR1 |
| ESR1 | ligand-dependent nuclear receptor | -0.2 | 0.0189 | AP2S1,ASS1,ATP6V1H,CA2,KPNB1,KRT17,MADD,NRCAM,POR,RAB5C,  RPL18A,SFPQ,SPAG9,TJP1,TJP2,TNPO1 |
| 1,2-dithiole-3-thione | chemical reagent | -0.2 | 0.0019 | DDX39B,PSMD1,PSMD5,RPL18,Snrpa (includes others),TXN |
| TP53 | transcription regulator | -0.2 | 1.68E-11 | ABAT,ACADM,ALDH1A1,ANXA1,ASS1,ATP5PB,CLASP1,CLTA,CRYAB,DCTN2,  DDX3X,DNAJA2,Dync1i2,ENO3,ETFA,EZR,FUBP1,GPHN,GPM6A,HADH,HK2,  HNRNPA2B1,KPNB1,LASP1,MYH9,PAICS,PLXNB2,PPM1A,PRKAR2A,PSMD1,  RAB5A,RAB5C,RPS18,RPS20,RPS25,RPS26,SFPQ,SYN1,TJP1,TOP2B,TRIM28,TTN |
| INSULIN (family) | group | -0.2 | 0.0447 | ACADM,CRYAB,DNAJA2,ETFA,GDI2,HK2,TCP1,TF,TJP1 |
| oleic acid | chemical - endogenous mammalian | -0.2 | 0.0498 | ABAT,ACADM,HADH,HK2 |
| tretinoin | chemical drug | -0.1 | 0.0112 | ACTR3,ALDH1A1,ALDH3A2,ANXA1,CA2,EEF1D,GNAO1,MYH9,NACA,NCL,POR,  RPL11,RPL3,RPS20,RPS4Y1,SYN1,TF,TJP1,TOP2B,TPT1 |
| RAF1 | kinase | -0.1 | 0.0024 | ANXA1,CA2,CRYAB,HNRNPA2B1,HNRNPAB,PLEC |
| PLX5622 | chemical drug | -0.1 | 0.00686 | ARF5,MAP1A,NCL,SNCB,SYN1,TJP1 |
| beta-estradiol | chemical - endogenous mammalian | 0.0 | 0.00055 | ABAT,ACADM,ALDH3A2,ANXA1,ARF5,ASS1,ATP5PB,CA2,CRYAA/LOC102724652,  CRYZ,EZR,FSCN1,GDI2,GTF2I,HADH,IPO5,KRT17,MGARP,MYH9,RAB5C,RGS9,  RPL14,RPL3,RPL8,RPS4Y1,SLC1A3,SLK,SPAG9,TF,TJP1,TJP2,TTN,TXN |
| HNF4A | transcription regulator | 0.0 | 0.000521 | ACTR3,ALDH1A1,ALDH5A1,ATP6V1H,CLTA,CRYZ,DDX39B,DNAJA2,GATD3/  LOC102724023,GTF2I,KPNB1,KRT17,NBEA,POR,PPME1,PSMD1,RAB10,RPL18,  RPL18A,RPS18,RPS20,RPS25,SF3B1,SUMO2,TF,TXN,UBE2V1 |
| phorbol esters | chemical - other | 0.0 | 0.00253 | ANXA1,CRYAA/LOC102724652,HK2,TXN |
| mir-210 | microRNA | 0.0 | 0.00341 | ALDH5A1,LASP1,SMCHD1,TNPO1 |
| COPS5 | transcription regulator | 0.0 | 0.00408 | HNRNPU,KPNB1,NCL,PAICS,PLXNB2,RPL18 |
| pirinixic acid | chemical toxicant | 0.0 | 0.0228 | ACADM,ALDH3A2,HADH,HSD17B10,PAICS,POR,TXN |
| EGFR | kinase | 0.1 | 0.0296 | ANXA1,CRYAB,HK2,HNRNPA2B1,KPNB1,KRT17,NCL,PLXNB2 |
| ESRRA | transcription regulator | 0.1 | 0.046 | ACADM,ANXA1,HK2,MYH9,TTN |
| PPARA | ligand-dependent nuclear receptor | 0.1 | 0.0121 | ACADM,ALDH3A2,ASS1,HADH,HSD17B10,POR,TJP1,TXN |
| deoxycholate | chemical - endogenous mammalian | 0.2 | 0.00131 | ALDH1A1,HK2,SLC1A3,TJP1 |
| tamoxifen | chemical drug | 0.2 | 0.0269 | ALDH3A2,ASS1,CA2,CRYZ,NRCAM,TXN |
| IL15 | cytokine | 0.2 | 0.035 | ANXA1,ENO3,GDI2,HK2,HNRNPA2B1,TJP1,TJP2 |
| TGFB2 | growth factor | 0.2 | 0.0392 | ALDH1A1,CA2,HK2,SLC6A11 |
| PPARGC1A | transcription regulator | 0.2 | 0.0383 | ABAT,ACADM,ALDH5A1,GNAO1,HK2,OPN1LW,PACSIN2 |
| puromycin aminonucleoside | chemical reagent | 0.2 | 0.00479 | ACTR3,ANXA1,ETFA,HADH |
| KRAS | enzyme | 0.2 | 0.0351 | ACTR3,CRYAB,CRYZ,CTNND2,GTF2I,HK2,KPNB1,MADD,NCL,PI4KA,POR,TCP1 |
| metribolone | chemical reagent | 0.2 | 0.000365 | ATP5PB,ATP6V1E1,CRYAB,CRYZ,CTNND2,ETFA,EZR,HADH,HK2,HSD17B10,POR |
| CD3 (complex) | complex | 0.3 | 0.00211 | ACTR3,ANXA1,CLIP1,GDI2,HNRNPA2B1,HUWE1,ILF2,NCL,PLEC,RPL30,UBE2V1 |
| TCR (complex) | complex | 0.3 | 0.00956 | ATP5PB,HADH,HSD17B10,RPL10A,RPL18A,RPL3,RPL30 |
| nicotine | chemical drug | 0.4 | 0.0179 | CA2,Gm21596/Hmgb1,GNAO1,ILF2,RAP1B |
| MYCL | transcription regulator | 0.4 | 0.000192 | DDX3X,GPHN,RPL10A,RPL11,RPL18 |
| E. coli B4 lipopolysaccharide | chemical toxicant | 0.4 | 0.0325 | ANXA1,HK2,RAB10,RPS25,TF |
| CIP2A | other | 0.4 | 0.0000827 | CRYAB,ENO3,HK2,LASP1,NCL |
| GRIN2A | ion channel | 0.4 | 0.00025 | GNAO1,KPNB1,PLXNB2,PSPC1,SFPQ |
| HBEGF | growth factor | 0.4 | 0.000485 | GNAT2,GNGT1,HK2,MYH9,OPN1LW |
| SNCA | enzyme | 0.4 | 0.01 | ALDH1A1,DCTN2,DMXL2,Ewsr1,EZR,SLC6A11,SYN1,VAT1L |
| SP2509 | chemical reagent | 0.4 | 0.044 | DDX3X,EZR,FUBP1,HNRNPA2B1,KPNB1 |
| MTOR | kinase | 0.5 | 0.000112 | ACADM,ATP5PB,ETFA,GPHN,HK2,MADD,PGP,PRKAR2A,RPS18,UBE2O |
| geldanamycin | chemical drug | 0.5 | 0.000417 | ANXA1,CACYBP,DDX3X,GNAO1,PPM1A,RAB10,RAB5A,RAB5C |
| IL4 | cytokine | 0.5 | 0.0152 | ANXA1,ASS1,CA2,CHD4,CLIP1,FSCN1,HK2,MYH9,NCL,PLEC,PRRC2C,  RAB5A,RELCH,Snrpa (includes others),TJP2 |
| FOS | transcription regulator | 0.6 | 0.00434 | CA2,CLIP1,DDX3X,EZR,HK2,HSD17B10,MYH9,RPS18,SUMO2,TXN |
| QKI | other | 0.6 | 0.0000687 | CLTA,FSCN1,GPHN,GPM6A,RAB5A,RAB5C |
| NFKBIA | transcription regulator | 0.6 | 0.00796 | CRYAB,DDX3X,FSCN1,GATD3/LOC102724023,Gm21596/Hmgb1,HK2,RPL8,RPS18 |
| hydrogen peroxide | chemical - endogenous mammalian | 0.6 | 0.0407 | ANXA1,CRYAB,DNAJA2,EZR,PRKCSH,RPL7A,RTN3,TXN |
| KCNJ2 | ion channel | 0.7 | 0.00000103 | CLASP1,CLIP1,Ktn1,MAP1A,MYEF2,MYH14,MYH9,PLEC |
| GnRH analog | biologic drug | 0.7 | 0.00112 | ATP6V1E1,CLTA,GATD3/LOC102724023,GDI2,PI4KA,PSMD1,RAB5C,UBE2V1 |
| CD28 | transmembrane receptor | 0.7 | 0.00176 | ACTR3,ANXA1,CLIP1,HK2,HUWE1,ILF2,NCL,RPL30 |
| lipopolysaccharide | chemical drug | 0.8 | 0.00173 | ACADM,ALDH1A1,ANXA1,ASS1,CRYAB,DDX3X,EEF2,FSCN1,GDI2,Gm21596/  Hmgb1,GPM6A,HK2,IPO5,LASP1,MYH14,MYH9,NCL,NIPSNAP2,PI4KA,PPM1A,  RAB10,RAB5A,Snrpa (includes others),SPAG9,SUMO2,TF,TJP1,TPT1,TXN,UBE2V1 |
| FOXA1 | transcription regulator | 0.8 | 0.0182 | ANXA1,HADH,HK2,MYH9,TF |
| CX3CR1 | G-protein coupled receptor | 0.8 | 0.000885 | CLASP1,CRYAA/LOC102724652,CRYAB,SFPQ,SYN1,TF |
| forskolin | chemical toxicant | 0.8 | 0.0479 | CRYAA/LOC102724652,CRYAB,EZR,HK2,PRKAR2A,RAB5A,RAB5C,RPS20,TJP1 |
| TNF | cytokine | 0.9 | 0.0312 | ACADM,ANXA1,ASS1,CA2,CLASP1,CRYAB,FSCN1,GATD3/LOC102724023,  GNAT2,GNGT1,HK2,MYH9,OPN1LW,PLXNB2,RGS9,SLC1A3,TF,TJP1,TJP2,TPT1,TXN |
| HIF1A | transcription regulator | 0.9 | 0.0341 | ENO3,FSCN1,HK2,HUWE1,MYH9,TJP1,TTN,TXN |
| TSC2 | other | 0.9 | 0.021 | ANXA1,CRYAB,HK2,PSMD1 |
| gentamicin | chemical drug | 1.0 | 0.000359 | ABAT,ACADM,ALDH1A1,DNAJA2,GPM6A,HNRNPU,NCL,SFPQ,TJP1 |
| CPT1B | enzyme | 1.0 | 0.000371 | ACADM,ATP6V1H,CA2,FUBP1,HADH,HK2,PRKAR2A |
| mono-(2-ethylhexyl)phthalate | chemical toxicant | 1.0 | 0.0206 | ACADM,ATP5PB,HADH,HK2 |
| VEGFA | growth factor | 1.0 | 0.0486 | ACADM,ETFA,RAB5A,TJP1,TJP2 |
| ADIPOR1 | transmembrane receptor | 1.1 | 0.00233 | ACADM,Hmgb3,OPN1LW,PLEC |
| FGFR1 | kinase | 1.1 | 0.0195 | HK2,PLXNB2,SFPQ,TXN |
| lactic acid | chemical - endogenous mammalian | 1.1 | 0.00851 | CACYBP,CEND1,HK2,NRCAM,TXN |
| TGFB1 | growth factor | 1.1 | 0.00584 | ALDH3A2,ALDH5A1,ASS1,CHD4,ENO3,FSCN1,FUBP1,GATD3/LOC102724023,  GNAO1,HADH,HK2,HNRNPAB,HSD17B10,KRT17,LASP1,MYH9,NBEA,PPM1A,  PSMD1,RAB1A,TJP1,TJP2,VAT1L |
| BDNF | growth factor | 1.2 | 0.0196 | EEF1D,EEF2,FSCN1,GNAO1,PLXNB2,SYN1 |
| RCN3 | other | 1.3 | 0.0000154 | RPL11,RPL3,RPL8,RPS18,RPS25 |
| fluoride | chemical - endogenous mammalian | 1.3 | 0.000163 | ACTR3,DDX39B,GNAO1,HNRNPA2B1,TOP2B |
| RHO | G-protein coupled receptor | 1.4 | 0.00194 | ACADM,GNAT2,GNGT1,HK2,RGS9,SF3B1 |
| LH (complex) | complex | 1.6 | 5.98E-14 | ATP5IF1,EEF2,EZR,HK2,PRKAR2A,PSMD1,RAB1A,RAB5A,RAB5C,RPL10A,  RPL18A,RPL3,RPL30,RPL8,RPS18,RPS20,RPS25,RPS26,Rps3a1,RPS4Y1 |
| E2F1 | transcription regulator | 1.7 | 0.00721 | ACTR3,CA2,CRYAB,DDX39B,NCL,RAB1A,SMARCC2,TOP2B,TRIM28 |
| uranyl nitrate | chemical toxicant | 1.7 | 2.73E-08 | ATP5IF1,CRYZ,EEF2,GATD3/LOC102724023,HNRNPA2B1,RAB5C,RPL10A,  RPS26,TPT1 |
| miR-124-3p (and other miRNAs w/seed AAGGCAC) | mature microRNA | 2.0 | 0.0626 | HADH,MTPN,MYH9,TJP2 |
| HSP90B1 | other | 2.0 | 0.000513 | ARF5,RAB10,RPL30,RPS20 |
| LEP | growth factor | 2.0 | 0.219 | ACADM,ASS1,ATP5PB,Gm21596/Hmgb1,SYN1 |
| MYCN | transcription regulator | 2.0 | 8.95E-10 | ALDH1A1,EEF1D,EEF2,HK2,MYH9,NACA,NCL,RPL11,RPL18,RPL18A,RPL3,  RPL30,RPL8,RPS20,RPS25,RPS26 |
| 3,5-dihydroxyphenylglycine | chemical reagent | 2.2 | 3.2E-17 | ACTR3,ALDH1A1,RPL10A,RPL11,RPL14,RPL18,RPL18A,RPL3,RPL30,RPL7A,  RPL8,RPS18,RPS20,RPS25,RPS26,RPS4Y1,TPT1 |
| AGN194204 | chemical drug | 2.2 | 0.0133 | ASS1,CA2,ENO3,HK2,Hmgb3 |
| MLXIPL/ChREBP | transcription regulator | 2.3 | 4.09E-14 | RPL10A,RPL11,RPL14,RPL18,RPL18A,RPL3,RPL30,RPL7A,RPL8,RPS18,  RPS20,RPS25,RPS26,Rps3a1,RPS4Y1 |
| MYC | transcription regulator | 2.4 | 4.66E-14 | ACADM,ACTR3,ADD1,ASS1,ATP5IF1,CRYAB,DDX39B,DDX3X,EEF2,EZR,  GDI2,HK2,HNRNPA2B1,HNRNPAB,HNRNPU,KRT17,NCL,PAICS,RAB10,  RPL10A,RPL11,RPL14,RPL18,RPL18A,RPL3,RPL30,RPL7A,RPL8,RPS18,  RPS20,RPS25,RPS26,Rps3a1,RPS4Y1,SUMO2,TF,TXN |
